# Multiple Measures Reveal The Value of Both Race And Geographic Ancestry For Self-Identification

**DOI:** 10.1101/701698

**Authors:** Vincent Damotte, Chao Zhao, Chris Lin, Eric Williams, Yoram Louzoun, Abeer Madbouly, Rochelle Kotlarz, Marisa McDaniel, Paul J. Norman, Antoine Lizee, Natalie M. Myres, Catherine A. Ball, Kenneth G. Chahine, Jake Byrnes, Yong Wang, Martin Maiers, Jill A. Hollenbach

## Abstract

There is long-standing tension regarding whether and how to use race or geographic ancestry in biomedical research. We examined multiple self-reported measures of race and ancestry from a cohort of over 100,000 U.S. residents alongside genetic data. We found that these measures are often non-overlapping, and that no single self-reported measure alone provides a better fit to genetic ancestry than a combination including both race and geographic ancestry. We also found that patterns of reporting for race and ancestry appear to be influenced by participation in direct-to-consumer genetic ancestry testing. Our results demonstrate that there is a place for the language of both race and geographic ancestry as we seek to empower individuals to fully describe their family history in research and medicine.

**One Sentence Summary:** Self-identification in the United States according to both racial and geographic terms best reflects genetic ancestry in individuals.

## Main Text

“The guy taking the census came to my door and he was asking about my self-identification, so I said ‘Greek.’ And he said, okay, I’ll put ‘White.’ And then I said, no, not ‘White’, point to me on the map where it says ‘White’” ~overheard, San Francisco International Airport

How do we ensure inclusion of diverse populations for the next generation of genomic, biomarker, behavioral research, and clinical trials *(1)*? Historically, subject participants in biomedical research have provided self-identification using race categories as defined by the United States Office of Management and Budget (OMB) *(2)*; indeed, federally funded researchers are mandated to collect and report this information. However, in the interest of stepping away from the socially loaded and disputed concept of race, many researchers have sought to focus rather on identification according to geographic ancestry *(3–5)*. It is argued that these measures better reflect human history and are more likely to represent biological differences compared to race, which is often described as a social construct *(6)*. Nevertheless, there is limited consensus *(7)* and a long-standing tension *(8)* regarding whether and how to use both race and ancestry in biomedical research *(9)*. Thus, debates over the utility of race versus geographic ancestry in genomics and biomedical research continue unabated *(10)*, with some arguing the language of “race” should be abandoned completely *(11)*.

Previous work has demonstrated that measures of race *(12)* (how people describe themselves using official racial categories) as well as ancestry *(13)* (how people describe their family origins in terms of geographic locations) each serve as reasonable proxies for genetic ancestry. However, whether and how these measures intersect has not been examined in the context of genetic variation. Examining simultaneously, and making the distinction between, the specific dimensions of race and geographic ancestry is necessary, because each draw on distinct aspects of people’s identities. At the same time, both are commonly employed as a proxy for genetic ancestry and are often used interchangeably in biomedical research *(7)*. Race may be tied to individuals’ socioeconomic status, lived experiences of marginalization or discrimination, and group affiliation *(14)*. Self-reported ancestry reflects what people know about the geographic origins of their ancestors, which is closely tied to patterns of family socialization *(15)*.

Although previous investigations have examined the relationship between single measures of self-identification and genetic ancestry *(12, 13, 16, 17)*, here we expand on our earlier work *(18)* with an approach that differs from these studies in several important ways. We directly incorporate findings from the social sciences (*19*) to perform the first large-scale study comparing multiple measures of self-identification simultaneously with genetic ancestry in the same cohort. We leverage the genetic information to facilitate comparison between measures and understand whether some are more closely related to genetic ancestry than others. We do so in a large and diverse sample of the U.S. adult population (Tables S1 and S2), considering how both race and ancestry can be used to best describe human diversity.

We collected multiple self-reported measures of race and ancestry from a cohort of more than 100,000 U.S. adults who also provided genetic data as potential donors registered with the National Marrow Donor Program (NMDP). To ascertain genetic ancestry, we used the registry’s data for the human leukocyte antigen (HLA) complex on chromosome 6, which is critical to matching in tissue transplant. The HLA loci exhibit extreme levels of variability and differentiation among human populations, and thus can be used as ancestry informative markers *(20–23)*. Our survey of potential NMDP donors, conducted for this study in spring 2015, included questions about racial self-identification and multiple (geographic) ancestry items. For self-reported ancestry, we included three measurement approaches: 1) personal ancestry (PA), a check-all-that-apply option using a series of geographic categories; 2) personal ancestry salience (PAS) a measure that asked people to “weight” their ancestry self-reports on a 100-point scale; and 3) family ancestry (FA), check-all-that-apply ancestry questions about specific biological relatives, such as grandparents. In order to fully exploit the FA responses, we also computed a summary family fractional ancestry (FFA) value from the family responses based on the number of ancestry selections per parent or grandparent. In addition to asking respondents to describe themselves using official racial categories (RC), we also asked that they tell us how they think other Americans would classify them using the same categories, which we term “reflected race” (RR)(*24*). The complete survey is provided in File S1.

We found that measures of self-reported race and ancestry are often non-overlapping, even when administered simultaneously in the same cohort. On the surface, responses for RC and PA might seem to provide redundant information, with many respondents identifying as White and also reporting PA from Western Europe, for example. However, cross-tabulating the measures with one another showed they are not as interchangeable as they might appear at first glance. When comparing racial self-identification and PA, every possible PA was connected to every possible RC in our sample (Fig. 1A), yielding a total of 3,582 different RC/PA combinations (Table S3). Nearly 60% of the sample self-reported two or more PA responses, and close to 12% provided two or more RC responses. Even when we restrict to individuals who selected a single PA and single RC response to describe themselves (39% of our sample), much of the complexity between ancestry and race reporting remains (Fig. 1B).

**Fig. 1.**
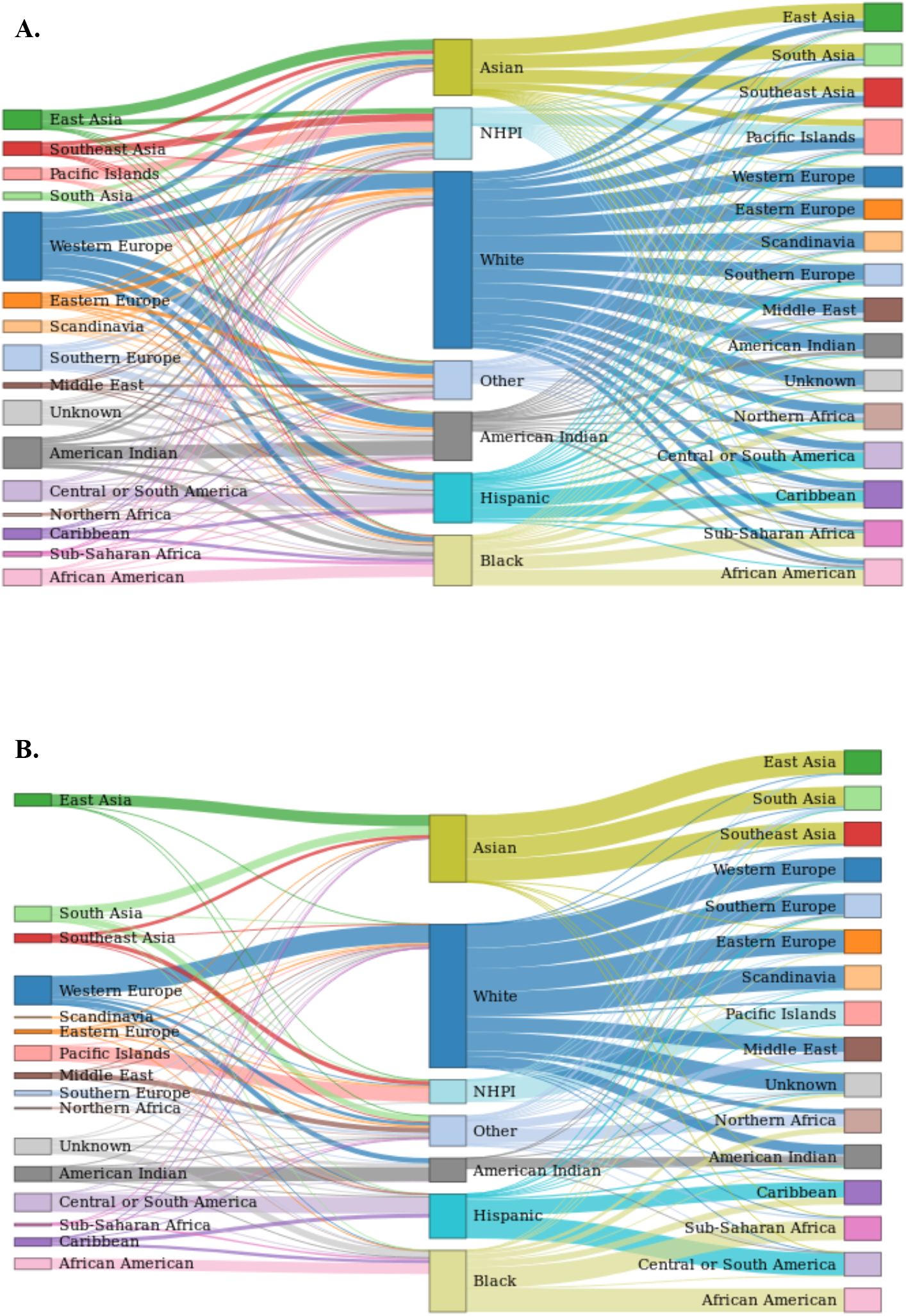
Sankey diagrams of connection between racial categories and geographic ancestries selected by respondents. (A) All respondents were considered, (B) only respondents who selected a single race category and a single geographic ancestry were considered.

To understand how these measures of self-reported race and ancestry relate to genetic ancestry, we employed a Bayesian classifier to assign the most probable geographic origin for subjects’ HLA haplotypes. Our previous work had shown that population-level HLA haplotype ancestry assignments using this method are equivalent to ancestry proportions derived from a well-characterized panel of ancestry informative markers *(18)*. To further validate the classifier, we examined prediction of the HLA-based ancestry classifications from ancestry proportions derived from over 700,000 single nucleotide polymorphism (SNP) markers for an independent dataset of 1,983 individuals, with cross-validation revealing accuracy approaching 85% (Supplemental Methods).

We tested fit of all self-reported race and ancestry responses alone and in specific combinations as predictors of genetic (HLA haplotype) ancestry in a multinomial logistic regression model, including covariates for age, sex, and educational attainment. Our survey methodology included randomly switching the order in which the race vs. ancestry sections were presented, which yielded some variation in the number of responses for each section, and thus we adjusted for this feature. Likewise, we adjusted for the email outreach recruiting participants, the specific language of which varied (Fig. S1). We found that no single self-reported measure of race or ancestry alone provides a better fit to genetic ancestry classification than combined measures (Fig. 2). When examining single measures, PA provided the best model fit, lending support to the notion that geographic ancestry serves as a better proxy for genetic ancestry than race. However, RC performed better than any of our other single measures, including FA, while RR performed very poorly, with the lowest R2 of any measure. Our quantitative measures, PAS and FFA, were highly correlated (Table S4), but had the highest misclassification rates of any single measure we examined, diminishing the overall model fit. Although PA performed better than the RC response alone, fit to genetic ancestry was significantly improved by incorporating the RC response with any of the ancestry measures, with the most significant improvements noted for combinations including PA and FA. Strikingly, the best-fitting model predicting genetic ancestry classification included a combination of RC self-identification and PA. This combined measure showed marked improvement in model fit compared to the PA single measure (p<0.001).

**Fig. 2.**
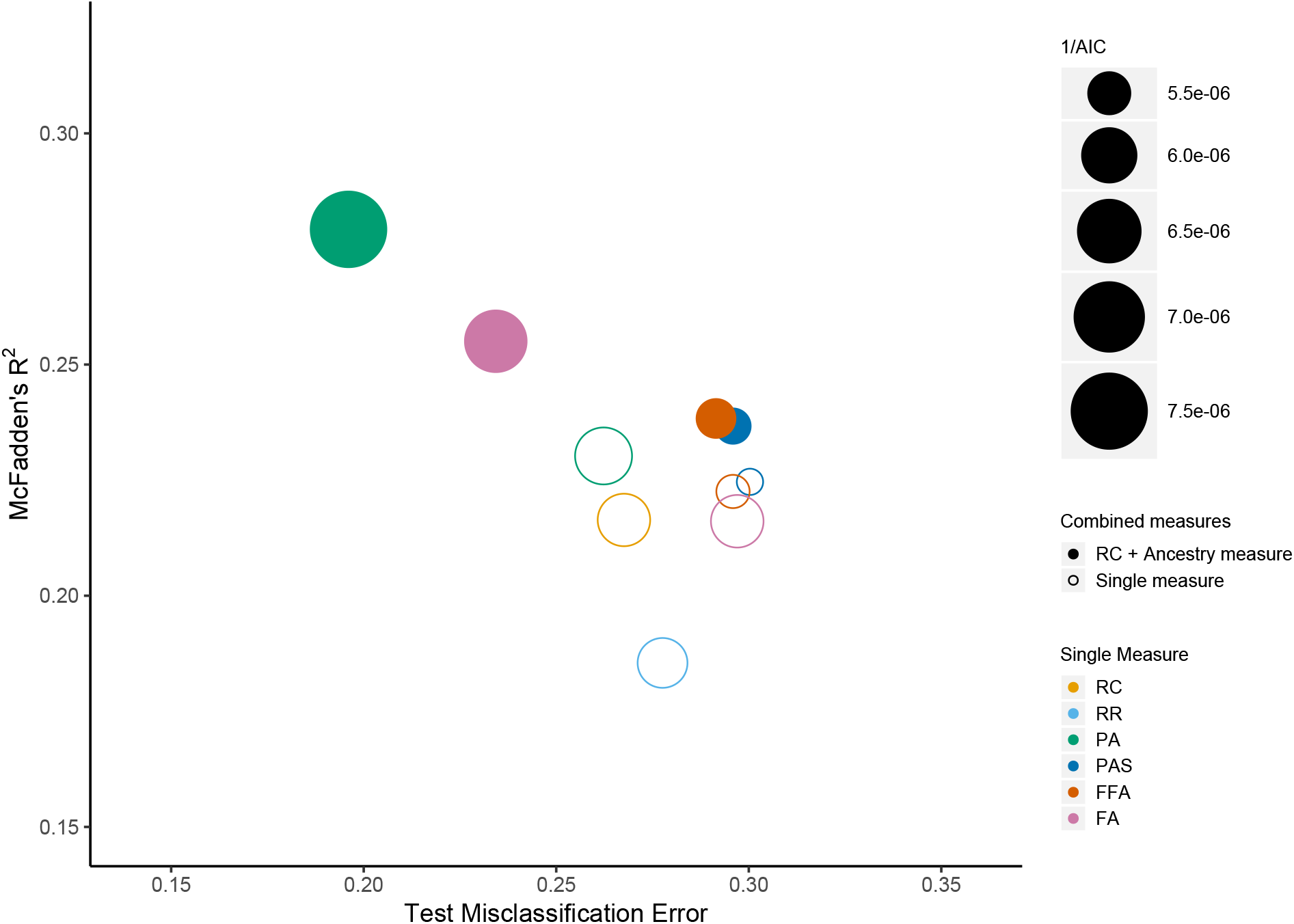
Performance assessment of different races and/or ancestries models. These models represent the performance of the fits of different models as predictors of genetic (HLA haplotype) ancestry (see Materials and Methods). RC: race category; PR: personal race; RR: Reflected race; PA: Personal Ancestry; PAS: Personal Ancestry Salience; FFA: Fractional family ancestry; FA: Family Ancestry

Specific examples from our data illustrate why combining race and ancestry responses serves to better represent genetic variation than single measures of self-identification. For instance, complexity in reporting American Indian race and ancestry is well documented in demographic studies *(25, 26)*. American Indian PA is reported frequently in our sample (15% of individuals), and is most often seen in combination with Western European PA (N=5709). Despite the fact that “American Indian” is also provided as an option for the RC response, many individuals reporting this PA combination report only the White RC. We computed the genetic distance between individuals reporting the specific combination of Western Europe and American Indian PA with only White RC (80%) and those who reported the same PA (Western Europe and American Indian) with White RC plus American Indian RC (17%) or only American Indian RC (1.6%); using a permutation procedure, we found that the White-only RC and White RC plus American Indian RC groups are not significantly divergent (p=0.15). However, the American Indian-only RC group is significantly divergent from the White-only RC group (p<0.001) and from the White RC plus American Indian RC group (p=0.03), showing the added value of combining race and ancestry responses.

Whereas incorporation of salience values (PAS) did not improve the overall fit of our models, they do provide important insights into the underlying dynamics in ancestry identification. Although frequently reported, American Indian PA yields the lowest mean PAS value (16.8) of any PA response (Fig. 3). Even among individuals who report American Indian FA for all four of their biological grandparents, their mean American Indian PAS value is only 49; in comparison, individuals who report four South Asian grandparents FA report South Asian mean PAS of 99 (p<0.001). These results may also explain why PA provided better overall model fit to genetics than FA. Notably, individuals who identify with American Indian RC report significantly higher American Indian PAS than those who did not (mean 26 vs. 14; p<0.001). In this case, racial self-identification appears to signal both personal and biological relevance for this particular ancestry. Our results also illustrate one of the pitfalls of using a check-all-that-apply format for reporting geographic origins as the sole self-identification measure in biomedical research.

**Fig. 3.**
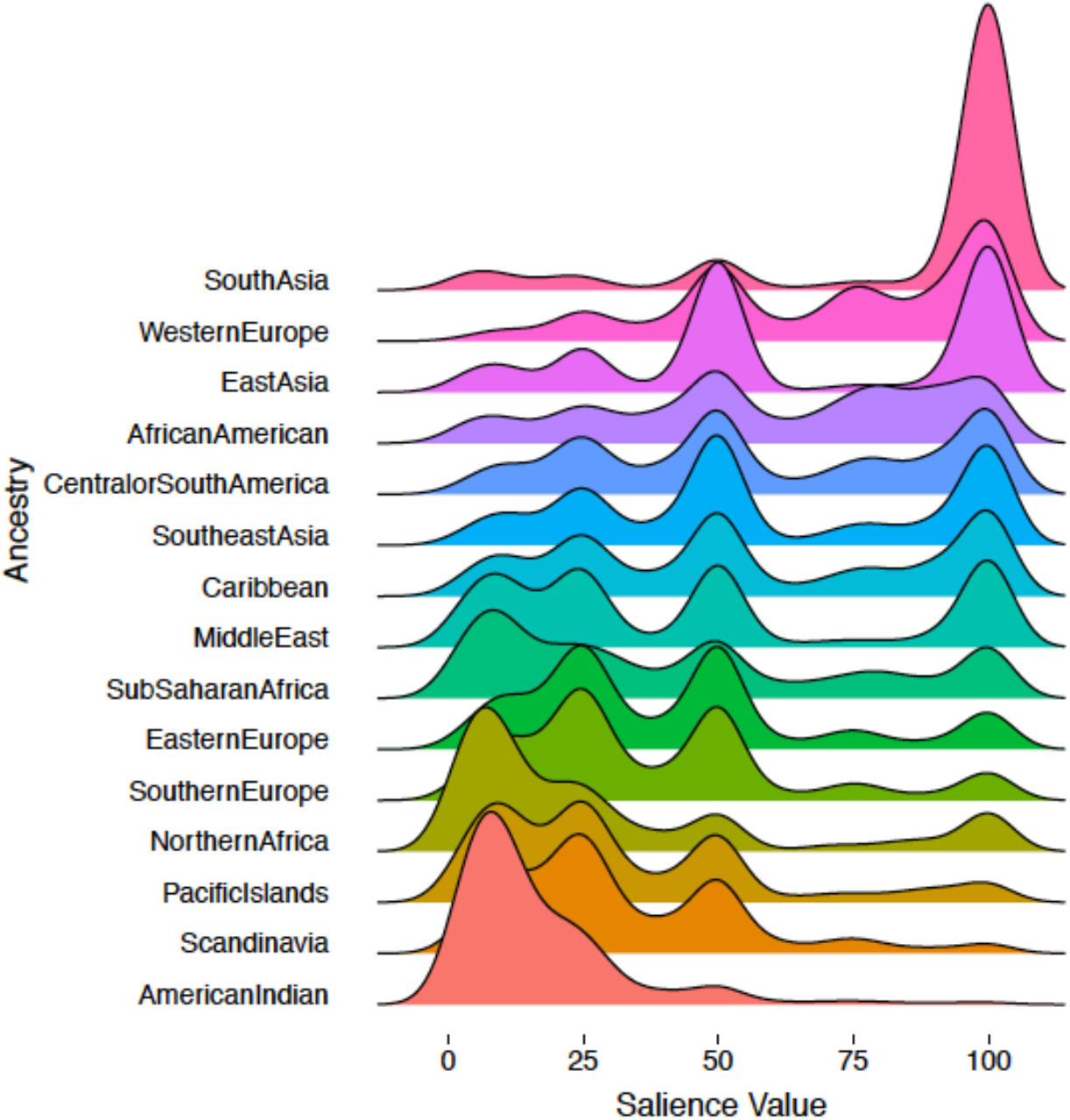
Density plots of personal ancestry salience (PAS) values given by individuals who selected specific geographic ancestry.

Likewise, we observed complexity comparing racial self-identification as Black with sub-Saharan African PA, furthering support for combining measures of race and ancestry when seeking to use self-identification as a surrogate for genetic variation. Although tracing ancestry to the original peoples of sub-Saharan Africa is the official definition of the “Black or African American” racial category in the U.S. *(2)*, many descendants of former slaves know little about their pre-slavery geographic origins *(27)*. To acknowledge this, we offered both “Sub-Saharan Africa” and “African American” categories among our ancestry responses. Among respondents who identified RC as Black alone (N=3038), 67% reported African American PA, compared to 17% who reported Sub-Saharan African PA. We analyzed the genetic distance between individuals who identified as Black RC alone and who reported African American ancestry only and those who reported Sub-Saharan African ancestry only and found significant divergence (p<0.001). One explanation for these observations may be found in respondents’ nativity: among respondents who identified as Black RC alone, respondents who reported sub-Saharan African ancestry were significantly less likely to have been born in the U.S. than those who did not report this ancestry (84% and 93% respectively; p<0.001). Foreign-born Black RC respondents who reported sub-Saharan African PA also reported a mean sub-Saharan African PAS value of 82, compared to 45 for their U.S.-born counterparts who selected the same RC and PA responses (p<0.001). Thus, although shared racial identification suggests a shared social experience of “blackness,” which likely has implications for health *(28, 29)*, a study that recruits subjects solely by racial self-identification might miss the genetic variation among individuals and their differing immigration histories, both of which could be important for understanding health disparities. For some other ancestries racial self-identification has even more limitation. A high proportion of individuals claiming only Middle Eastern or North African ancestry do not identify with any of the standard OMB RC’s, and rather select Other. Likewise, South Asian ancestry is generally split between the Other category and Asian RC (Fig. 1). These results underscore the notion that race and ancestry are describing distinct aspects of self-identification, which partially – but far from completely – overlap. Moreover, these patterns vary by population, emphasizing the need to embrace multiple measures in order to offer appropriate options to diverse cohorts.

Finally, we found that self-identification reporting patterns may be transformed by participation in direct-to-consumer genetic ancestry testing (GAT). Approximately 5% of our respondents reported having taken a GAT *(30)*. Overall, these individuals gave more responses for ancestry (mean responses 2.3 vs 1.9; p<0.001) as well as distinctive combinations of race and ancestry reporting compared to those who did not use GAT. Among respondents who identified as Black RC alone, 62% reported sub-Saharan African PA if they had taken a GAT compared to 14% who have never taken a GAT (p<0.001). In contrast, these groups reported African American PA nearly equivalently at 70% and 66%, respectively. In contrast to the larger sample, genetic distance measures were non-significant between Black RC individuals who did or did not report sub-Saharan African PA. Likewise, among GAT participants, 96% of Black respondents reporting sub-Saharan ancestry also reported being U.S. born. In addition to sub-Saharan African PA, a number of other PA responses were also found to differ in frequency according to whether respondents had used GAT. For example, among GAT takers, American Indian PA was reported less often by individuals identifying as White RC (pcorr=0.004), but more often by individuals identifying as Hispanic RC (pcorr<0.001) compared to those who did not use GAT. Thus, in contrast to individuals who did not participate in GAT, here the race response did not improve model fit and racial identification appears not relevant with respect to genetic ancestry.

Taken together, our results demonstrate that there is a place for the language of both race and (geographic) ancestry as we seek to empower individuals to fully describe their family history in research and medicine. They highlight the importance of using measures of both race and ancestry to allow for a range of processes, both social and biological, rather than contrasting measures in a way that assumes one is the more objectively correct classification. For example, it may be enticing to treat self-reported geographic ancestry as a better proxy for biology, as part of an attempt to sidestep politicized notions of “race” in biomedical settings. However, our results demonstrate that while providing important information, self-reported geographic ancestry alone is not as good a proxy for genetic variation as when coupled with racial self-identification, and there is ample research that shows self-reported race has a role to play in studies of health disparities. Racial distinctions among humans were not scientifically valid to begin with, but their use in contemporary research helps to acknowledge and account for the differing social experiences of people living in a country with a long history of racial stratification. Our results for individuals participating in GAT suggest that as genealogical tools and technologies increase in popularity and accessibility, individuals may move toward means of self-identification that are more geographically, and less racially, based. Meanwhile, we stand to gain better understanding of patterns of health and illness by recognizing the differences between measures of race and ancestry, and leveraging instances of empirical convergence and divergence for insight into biological processes.

## Acknowledgments

The authors wish to thank Aliya Saperstein for assistance in the conception and design of the survey. We also thank the study participants.

## Funding

This work was supported by grants from NIH National Human Genome Research Institute (R21HG00804) and the Department of the Navy, Office of Naval Research (N00014-17-1-2388);

## Author contributions

Conceptualization: MM, JAH, RK, MM, EW; Data curation: VD, CZ, EW, MM Formal analysis: VD, CZ, CL, YL, AM, AL, NM, CB, KC, JB, YW, MM, JAH; Writing: VD, PJN, MM, JAH;

## Competing interests

Authors declare no competing interests;

## Data and materials availability

Analytical codes for this study are available at https://github.com/Hollenbach-lab/AQP_Paper1_PublicRelease. Request for data access must be sent to the corresponding author.

## Supplementary Materials

Materials and Methods

Figure S1

Tables S1-S4

References (*31-44*)

