## Supplementary Materials for "Multiple Measures Reveal The Value of Both Race And Geographic Ancestry For Self-Identification"

**This PDF file includes:**

Materials and Methods
Supplementary Text
Fig. S1
Tables S1 to S4

**Other Supplementary Materials for this manuscript include the following:**

File S1.

### Materials and Methods

Analytical codes for this study are available at:

https://github.com/Hollenbach-lab/AQP_Paper1_PublicRelease

Request for data access must be sent to the corresponding author.

Data cleaning

- Race data: Individuals with a non-available observation in at least one variable were removed.
- Personal Ancestry data: Individuals with a non-available observation in at least one variable were removed.
- Reflected Race data: Individuals with a non-available observation in at least one variable were removed.
- Personal Ancestry Salience data: Individuals with a non-available observation in at least one variable were removed. Only individuals with a sum of all salience different from zero were kept.
- Family Ancestry data: Family data were divided into two types of data: Individuals who selected ancestry data for their four grandparents and, when ancestry data were missing for at least one grandparent, we used the ancestry information selected for their two parents. Individuals with a non-available observation in at least one variable for the ancestry for their four grandparents or in at least one variable for the ancestry of their two parents were removed.

  First, per ancestry and per grandparent, the ancestry binary variables were transformed into percentages, by multiplying the binary outcome by 0.25. Then the same ancestry variables of each grandparent were summed up. Second, for individuals without ancestries for their four grandparents and with ancestries for at least one of their parents, ancestry variables were computed based on their parents ancestry information. Per ancestry and per parent, the ancestry binary variables were transformed into percentages, by multiplying the binary outcome by 0.50. If one of the parents did not have any ancestry information, we considered a percentage of 0.5 for the *Unknown* ancestry and 0 for the other choices for this parent. Next, the same ancestry variables of each parent were summed and the parents and grandparents ancestry percentages were bound to create the fractional family ancestry (FFA) data.

Descriptive analyses

The Sankey diagrams were made with the function *sankeyNetwork* in the R package networkD3 *(31)*. All the other plots were made with the R package ggplot2 *(32)*. Only ancestries weighted strictly above 0% were included for the ridgeline PAS analysis. Comparisons of PAS values between two groups were performed using a t-test. Comparisons of group size were performed using a Fisher test. Pearson's correlation coefficients were computed, and the correlation coefficients were tested for significance using Fisher's Z transformation.

Bayesian classifier to assign the most probable geographic origin for subjects' HLA haplotypes

The subjects were typed using a variety of PCR based DNA sequencing method targeting of HLA-A, -B, -C, -DRB1 and -DQB1 *(33)*. Subject were typed at 3 different laboratories from 2008 – 2014 during a period where the laboratory achieved an average unphased typing resolution score *(34)* of at least 0.75. The resulting genotypes have no phase information between loci but only modest amounts of ambiguity in terms of allele assignments within a locus. The genotypes were analyzed using an updated "multi-race" version of an imputation algorithm described previously *(35)* that generated: a list of all possible pairs of 5-locus phased, allele-level haplotypes with corresponding haplotype frequency and population origin for each, including cases where the two populations can either be the same or different.

The population frequencies used for this study were an updated version of those published previously *(36)*. 18 populations sets were included in the imputation process which are the 21 categories below excluding the three smallest groups (AISC, ALANAM, SCAMB).

| Race code | Detailed race/ethnic description |
|---|---|
| AAFA | African American |
| AFB | African |
| AINDI | South Asian Indian |
| AISC | American Indian – South or Central Am. |
| ALANAM | Alaska native or Aleut |
| AMIND | North American Indian |
| CARB | Caribbean black |
| CARHIS | Caribbean hispanic |
| CARIBI | Caribbean Indian |
| EURCAU | European caucasian |
| FILII | Filipino |
| HAWI | Hawaiian or other Pacific Islander |
| JAPI | Japanese |
| KORI | Korean |
| MENAFC | Middle Eastern or N. Coast of Africa |
| MSWHIS | Mexican or Chicano |
| NCHI | Chinese |
| SCAHIS | Hispanic – South or Central American |
| SCAMB | Black – South or Central American |
| SCSEAI | Southeast Asian |
| VIET | Vietnamese |

Estimate of Haplotype SIRE combination based on estimated single SIRE haplotype frequency distributions.

Given the possible genotypes of an individual *i* and a set of haplotype frequencies for each sub population – *k*, and haplotype $h_j$: $f_k(h_j)$, we first estimate the probability of each haplotype pair in each population pair $f_k(h_{j1}), f_l(h_{j2})$, where $h_{j1}, h_{j2}$ are a pair of haplotypes consistent with the genotype of individual *i.* We then estimate the total probability that individual *i* haplotypes originated from the population pair $k,l$ - $p(k,l) = \sum_{j1,j2} f_k(h_{j1}), f_l(h_{j2})$, where the sum is only on haplotypes consistent with the genotype of individual *i.* The race combination maximizing $p(k,l)$ is defined as the race combination of individual *i*. Given the race combination, we estimate the most probable haplotype on race k, as $h_1 = ArgMax(f_k(h_{j1}))$. The second haplotype is then defined as $h_2 = \underset{h_{j2}}{ArgMax}(f_l(h_{j2}) * f_k(h_1))$.

Estimation of ancestry proportions in validation sample

Admixture proportions were estimated for all 2,005 samples using ADMIXTURE (37). Each sample was analyzed separately in ADMIXTURE's "supervised" mode (--supervised) by comparing its genotypes with a pre-curated population reference panel consisting of 3,000 labeled reference samples from 26 global populations. The default block relaxation optimization method was utilized and the convergence criterion was set to a change in log-likelihood between two iterations falling below 0.01 (-C 0.01). In addition, standard errors were estimated for the admixture proportions by running ADMIXTURE with 40 bootstrap replicates (-B 40).

Population reference panel candidates were selected from the publicly accessible Human Genome Diversity Project (38, 39), an internal proprietary AncestryDNA reference collection, and a reference collection corresponding to AncestryDNA customers who explicitly provided prior consent to participate in research and have all family lineages tracing back to the same geographic region. All the candidates were analyzed through a quality control pipeline to remove samples with lower genotype call rate, samples genetically related to each other, and samples who appear as outliers from their purported population of origin based on Principal Component Analysis (PCA). Note, PCA analyses were performed using a subset of independent SNPs following linkage disequilibrium (LD) filtering. Using PLINK (40), one of each pair of high LD SNPs in a 50-SNP sliding window were removed. The final ethnicity reference panel contains 3,000 samples representing 26 distinct global populations. For more details, please see AncestryDNA's methods white paper (41).

Bayes classifier cross-validation

We used multinomial logistic regression to predict population assignments derived from the Bayes classifier from ancestry proportions derived from SNP data (as described above) in our validation cohort. The models were assessed by 10-fold cross-validation. The sample (N = 1981) was randomly partitioned into 10 approximately equal size subsamples. A single subsample is retained as the validation data for testing the model, and the remaining 9 subsamples are used as training data. The predictive accuracy was measured by root-mean-square error (RMSE) on both training data and test data. Our results indicate a testing misclassification error of only 16%, indicating very good agreement between the two methods of ancestry determination.

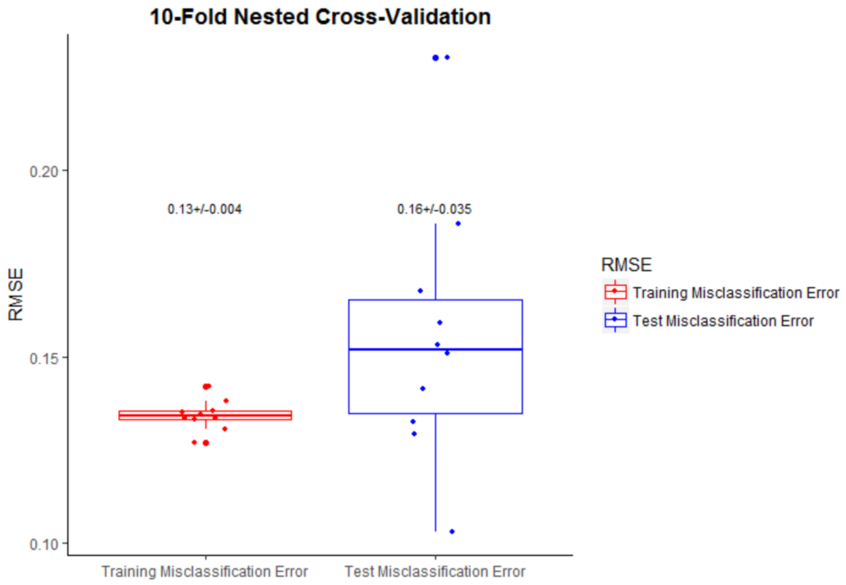


<u>Haplotypes clustering-based modeling</u>

1. *Races and Ancestries data preparation*

General cleaning previously performed was used to prepare the following survey data.

- Race (RC): No additional cleaning was done. Binary data for each variable, 7 variables.
- Personal Ancestry (PA): Individuals who only chose *Unknown* ancestry were removed. Binary data for each variable, 16 variables.
- Race + Personal Ancestry (RC/PA): The Race and Personal Ancestry information created above were combined. Binary data for each variable, 23 variables.
- Reflected Race (RR): The Reflected Race variable with 7 levels was converted into 7 variables: American Indian or Alaska Native, Asian, Black or African American, Hispanic or Latino, Native Hawaiian or other Pacific Islander, White and Other. Binary data for each variable, 7 variables.
- Personal Ancestry Salience (PAS): The Personal Ancestry Salience was used. No additional cleaning was done. Percentages data for each variable, 16 variables.
- Race + Personal Ancestry Salience (RC/PAS): The Race and Personal Ancestry Salience information created above were combined. Binary data for Personal Race variables and Percentages data for each Personal Ancestry Salience variable, 23 variables.

- Fractional Family Ancestry (FFA): Percentages (0 to 100%) data for each variable as prepared previously (see 1.Methods for data cleaning), 16 variables.
- Race + Fractional Family Ancestry (RC/FFA): The Race and Fractional Family Ancestry information created above were combined. Binary data for Races variables, Percentages (0 to 100%) data for Fractional Family Ancestry variables, 23 variables.
- Family Ancestry (FA): Above Fractional Familial Ancestry percentages (FFA) were replaced by 1 or 0, whether they were strictly higher than 0% or not. Binary data for each variable, 16 variables.
- Race + Family Ancestry (PR/FA): Race and Family Ancestry information created above were combined. Binary data for each variable, 23 variables.

*Commons IDs:* Only individuals present in each of these models were kept for subsequent analyses.

2. *K-means Clustering Algorithm*

Among 90,731 participants, there were 171 different unique haplotypes combinations based on 18 haplotypes. The unsupervised k-means clustering algorithm was performed to regroup and reduce the haplotypes levels for classification. The number of clusters was validated by the elbow method which run k-means clustering on the dataset for a range of values of k from 1 to 50, and the sum of squared errors (SSE) were calculated for each value of k. A smaller k value (k = 18) with a lower SSE was determined from the total within-cluster sum of squares plot (wss-plot) and was used to replace the 171 different unique genetic outcomes. The 18 initial "means" were randomly generated within the data domains. Each data point was assigned to its closest cluster center according to the Euclidean distance function. The new centroid or mean of all objects in each cluster were calculated. Each data point was reassigned to the new centroid. The steps were repeated until convergence has been reached.

3. *Multinomial Logistic Regression*

Multinomial logistic regression (MLR) was used to investigate the relationship between the population assignments based on HLA with race and ancestry information among 90,731 participants. MLR is designed to deal with cases of dependent variables with multiple classes.

One major advantage of MLR is it is robust to violations of assumptions of multivariate normality as is the case where there are some zero/one variables or where distributions are highly skewed or heavy-tailed. The strength of the MLR relationship between dependent variable and independent variables was estimated by correlation measure (pseudo R squares measures, such as McFadden's R2). Values of McFadden's $R^2$ between 0.2 and 0.4 are considered highly satisfactory of goodness of fit. To assess the strength of MLR relationship, the evaluation of the usefulness for logistic models was also considered. We assessed misclassification errors on both training set (81,657 participants) and test set (9,074 participants), which compared the predicted groups to the actual groups. MLR was performed using the R package *nnet (42)*.

Methods for Edward's Genetic Distances analyses

2,118 SNPs genotypes from the HLA region were available for 102,982 surveyed individuals. 443 SNPs were polymorphic and kept for subsequent analysis.

The adegenet R package *(43, 44)* was used to compute Edward's genetic distances. Only populations with more than 50 individuals were kept for analyses. 999 permutations were performed to obtain one-tailed p-values testing the genetic distances differences between two populations. A p-value below 0.05 signifies that no more than 49/999 permutations led to distances higher than the one observed.

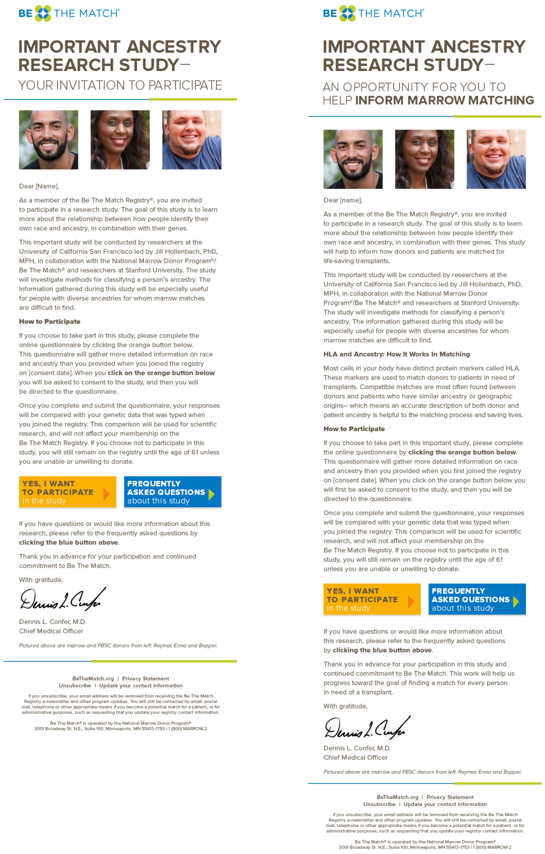

BE THE MATCH®

**IMPORTANT ANCESTRY RESEARCH STUDY–**

YOUR INVITATION TO PARTICIPATE

Dear [Name],

As a member of the Be The Match Registry®, you are invited to participate in a research study. The goal of this study is to learn more about the relationship between how people identify their own race and ancestry, in combination with their genes.

This important study will be conducted by researchers at the University of California San Francisco led by Jill Hollenbach, PhD, MPH, in collaboration with the National Marrow Donor Program®/Be The Match® and researchers at Stanford University. The study will investigate methods for classifying a person's ancestry. The information gathered during this study will be especially useful for people with diverse ancestries for whom marrow matches are difficult to find.

**How to Participate**

If you choose to take part in this study, please complete the online questionnaire by clicking the orange button below. This questionnaire will gather more detailed information on race and ancestry than you provided when you joined the registry on [consent date]. When you **click on the orange button below** you will be asked to consent to the study, and then you will be directed to the questionnaire.

Once you complete and submit the questionnaire, your responses will be compared with your genetic data that was typed when you joined the registry. This comparison will be used for scientific research, and will not affect your membership on the Be The Match Registry. If you choose not to participate in this study, you will still remain on the registry until the age of 61 unless you are unable or unwilling to donate.

**YES, I WANT TO PARTICIPATE** in the study | **FREQUENTLY ASKED QUESTIONS** about this study

If you have questions or would like more information about this research, please refer to the frequently asked questions by **clicking the blue button above**.

Thank you in advance for your participation and continued commitment to Be The Match.

With gratitude,

Dennis L. Confer, M.D.
Chief Medical Officer

*Pictured above are marrow and PBSC donors from left: Roymar, Erma and Bopper.*

BeTheMatch.org | Privacy Statement
Unsubscribe | Update your contact information

If you unsubscribe, your email address will be removed from receiving the Be The Match Registry e-newsletter and other program updates. You will still be contacted by email, postal mail, telephone or other appropriate means if you become a potential match for a patient, or for administrative purposes, such as requesting that you update your registry contact information.

Be The Match® is operated by the National Marrow Donor Program®
3001 Broadway St. N.E., Suite 100, Minneapolis, MN 55413-1753 | 1 (800) MARROW-2

BE THE MATCH®

**IMPORTANT ANCESTRY RESEARCH STUDY–**

AN OPPORTUNITY FOR YOU TO HELP **INFORM MARROW MATCHING**

Dear [name],

As a member of the Be The Match Registry®, you are invited to participate in a research study. The goal of this study is to learn more about the relationship between how people identify their own race and ancestry, in combination with their genes. This study will help to inform how donors and patients are matched for life-saving transplants.

This important study will be conducted by researchers at the University of California San Francisco led by Jill Hollenbach, PhD, MPH, in collaboration with the National Marrow Donor Program®/Be The Match® and researchers at Stanford University. The study will investigate methods for classifying a person's ancestry. The information gathered during this study will be especially useful for people with diverse ancestries for whom marrow matches are difficult to find.

**HLA and Ancestry: How It Works In Matching**

Most cells in your body have distinct protein markers called HLA. These markers are used to match donors to patients in need of transplants. Compatible matches are most often found between donors and patients who have similar ancestry or geographic origins– which means an accurate description of both donor and patient ancestry is helpful to the matching process and saving lives.

**How to Participate**

If you choose to take part in this important study, please complete the online questionnaire by **clicking the orange button below**. This questionnaire will gather more detailed information on race and ancestry than you provided when you first joined the registry on [consent date]. When you click on the orange button below you will first be asked to consent to the study, and then you will be directed to the questionnaire.

Once you complete and submit the questionnaire, your responses will be compared with your genetic data that was typed when you joined the registry. This comparison will be used for scientific research, and will not affect your membership on the Be The Match Registry. If you choose not to participate in this study, you will still remain on the registry until the age of 61 unless you are unable or unwilling to donate.

**YES, I WANT TO PARTICIPATE** in the study | **FREQUENTLY ASKED QUESTIONS** about this study

If you have questions or would like more information about this research, please refer to the frequently asked questions by **clicking the blue button above**.

Thank you in advance for your participation in this study and continued commitment to Be The Match. This work will help us progress toward the goal of finding a match for every person in need of a transplant.

With gratitude,

Dennis L. Confer, M.D.
Chief Medical Officer

*Pictured above are marrow and PBSC donors from left: Roymar, Erma and Bopper.*

BeTheMatch.org | Privacy Statement
Unsubscribe | Update your contact information

If you unsubscribe, your email address will be removed from receiving the Be The Match Registry e-newsletter and other program updates. You will still be contacted by email, postal mail, telephone or other appropriate means if you become a potential match for a patient, or for administrative purposes, such as requesting that you update your registry contact information.

Be The Match® is operated by the National Marrow Donor Program®
3001 Broadway St. N.E., Suite 100, Minneapolis, MN 55413-1753 | 1 (800) MARROW-2

**Fig. S1. The two different types of email outreach for recruiting participants.**

**Table S1. Survey respondents demographics (gender and age groups) separated by place of birth.**

Gender and age group were missing for 1 and 63 individuals, respectively. Place of birth was missing for 91 individuals, parents place of birth was missing for 39 individuals. Grand-parents place of birth was missing for 235 individuals.

| | **Total** | **Female** | **Male** | **[18-24]** | **[25-34]** | **[35-44]** | **[45-54]** | **[55-64]** | **[65+]** |
|---|---|---|---|---|---|---|---|---|---|
| **All respondents** | 103348 | 82226 | 21121 | 13576 | 35487 | 27804 | 18105 | 8313 | 9 |
| **RESPONDENT US BORN** | | | | | | | | | |
| Yes | 95770 | 80.1 % | 19.9 % | 13.3 % | 34.5 % | 26.6 % | 17.4 % | 8.2 % | 0 % |
| No | 7487 | 72.7 % | 27.3 % | 11.4 % | 32.8 % | 31.1 % | 18.5 % | 6.1 % | 0 % |
| **PARENTS US BORN** | | | | | | | | | |
| Neither | 11107 | 73.8 % | 26.2 % | 16.7 % | 35.6 % | 27.4 % | 15.4 % | 4.9 % | 0 % |
| One | 8190 | 80.7 % | 19.3 % | 17.2 % | 35.9 % | 26.2 % | 14.5 % | 6.2 % | 0 % |
| Both | 84012 | 80.2 % | 19.8 % | 12.3 % | 34.0 % | 26.9 % | 18.1 % | 8.6 % | 0 % |
| **GRANDPARENTS US BORN** | | | | | | | | | |
| None | 15319 | 74.7 % | 25.3 % | 14.8 % | 31.7 % | 25.6 % | 18.7 % | 9.1 % | 0 % |
| One | 3031 | 79.8 % | 20.2 % | 12.9 % | 27.6 % | 25.1 % | 22.3 % | 12.1 % | 0 % |
| Two | 13639 | 80.6 % | 19.4 % | 13.6 % | 31.7 % | 25.1 % | 19.5 % | 10.1 % | 0 % |
| Three | 9939 | 81.8 % | 18.2 % | 12.9 % | 34.8 % | 26.1 % | 17.7 % | 8.6 % | 0 % |
| Four | 61365 | 80.2 % | 19.8 % | 12.7 % | 35.9 % | 27.9 % | 16.5 % | 7.0 % | 0 % |

**Table S2. Survey respondents demographics (gender and age groups) separated by race.** Gender and age group were missing for 1 and 63 individuals, respectively.

| | **Total** | **Female** | **Male** | **[18-24]** | **[25-34]** | **[35-44]** | **[45-54]** | **[55-64]** | **[65+]** |
|---|---|---|---|---|---|---|---|---|---|
| **All respondents** | 103348 | 82226 | 21121 | 13576 | 35487 | 27804 | 18105 | 8313 | 9 |
| American Indian | 279 | 80.6 % | 19.4 % | 10.4 % | 22.9 % | 35.8 % | 24.0 % | 6.8 % | 0 % |
| Asian | 3461 | 67.6 % | 32.4 % | 18.2 % | 42.1 % | 25.1 % | 11.2 % | 3.4 % | 0 % |
| Black | 3044 | 84.3 % | 15.7 % | 14.1 % | 30.7 % | 29.1 % | 18.4 % | 7.7 % | 0 % |
| Hispanic | 4889 | 80.6 % | 19.4 % | 21.5 % | 34.9 % | 27.5 % | 12.4 % | 3.7 % | 0 % |
| Native Hawaiian or Pacific Islander | 129 | 76.7 % | 23.3 % | 10.1 % | 34.9 % | 36.4 % | 12.4 % | 6.2 % | 0 % |
| White | 78489 | 79.8 % | 20.2 % | 11.5 % | 33.4 % | 27.0 % | 18.9 % | 9.1 % | 0 % |
| Other | 1146 | 66.8 % | 33.2 % | 11.2 % | 34.9 % | 28.4 % | 17.0 % | 8.5 % | 0 % |
| Multi-race | 11903 | 81.4 % | 18.6 % | 18.9 % | 39.2 % | 25.4 % | 11.9 % | 4.5 % | 0 % |

**Table S3. (A) Number of races (rows) and ancestries (columns) selected. (B) Number of races (rows) and ancestries (columns) selected, considering only populations with more than 50 individuals.**

**A.**

| | **1** | **2** | **3** | **4** | **5** | **6** | **7** | **8** | **9** | **10** | **11** | **12** | **13** | **14** | **15** | **16** | **Total** |
|---|---|---|---|---|---|---|---|---|---|---|---|---|---|---|---|---|---|
| **1** | 40305 | 34549 | 13053 | 2810 | 524 | 102 | 38 | 12 | 1 | 3 | - | 1 | - | - | - | - | 91398 |
| **2** | 1224 | 4348 | 3211 | 1156 | 292 | 55 | 20 | 4 | 3 | - | 1 | - | - | - | - | - | 10314 |
| **3** | 88 | 206 | 542 | 345 | 132 | 32 | 15 | 5 | 2 | 1 | - | - | - | - | - | - | 1368 |
| **4** | 11 | 14 | 24 | 70 | 35 | 22 | 9 | 1 | 1 | 1 | - | - | - | - | - | - | 188 |
| **5** | 1 | 1 | 3 | 4 | 6 | 6 | 2 | - | 1 | 4 | - | - | - | - | 1 | - | 29 |
| **6** | - | - | - | - | 1 | - | - | - | 1 | - | - | - | - | - | - | - | 2 |
| **7** | - | - | - | - | - | - | - | - | - | - | - | - | - | - | - | - | - |
| **Total** | 41629 | 39118 | 16833 | 4385 | 990 | 217 | 84 | 22 | 9 | 9 | 1 | 1 | | | 1 | | 103299 |

**B.**

| | **1** | **2** | **3** | **4** | **5** | **6** | **7** | **8** | **9** | **10** | **11** | **12** | **13** | **14** | **15** | **16** | **Total** |
|---|---|---|---|---|---|---|---|---|---|---|---|---|---|---|---|---|---|
| **1** | 39913 | 33144 | 11307 | 1918 | 61 | - | - | - | - | - | - | - | - | - | - | - | 86343 |
| **2** | 622 | 3103 | 1845 | 253 | - | - | - | - | - | - | - | - | - | - | - | - | 5823 |
| **3** | - | - | 107 | - | - | - | - | - | - | - | - | - | - | - | - | - | 107 |
| **4** | - | - | - | - | - | - | - | - | - | - | - | - | - | - | - | - | - |
| **5** | - | - | - | - | - | - | - | - | - | - | - | - | - | - | - | - | - |
| **6** | - | - | - | - | - | - | - | - | - | - | - | - | - | - | - | - | - |
| **7** | - | - | - | - | - | - | - | - | - | - | - | - | - | - | - | - | - |
| **Total** | 40535 | 36247 | 13259 | 2171 | 61 | - | - | - | - | - | - | - | - | - | - | - | 92273 |

**Table S4. Correlation of fractional family ancestry (FFA) with personal ancestry salience (PAS).**

| Ancestry | Correlation | P-value |
|---|---|---|
| **South Asia** | 0.94 | < 0.001 |
| **East Asia** | 0.92 | < 0.001 |
| **Middle East** | 0.89 | < 0.001 |
| **Southeast Asia** | 0.83 | < 0.001 |
| **Eastern Europe** | 0.80 | < 0.001 |
| **Northern Europe** | 0.75 | < 0.001 |
| **Western Europe** | 0.74 | < 0.001 |
| **Southern Europe** | 0.73 | < 0.001 |
| **Caribbean** | 0.73 | < 0.001 |
| **Central or South America** | 0.72 | < 0.001 |
| **Scandinavia** | 0.69 | < 0.001 |
| **Pacific Islands** | 0.64 | < 0.001 |
| **Sub-Saharan Africa** | 0.62 | < 0.001 |
| **African-American** | 0.61 | < 0.001 |
| **American Indian** | 0.41 | < 0.001 |
