## Supplementary File S1 for "Multiple Measures Reveal The Value of Both Race And Geographic Ancestry For Self-Identification"

### Consent form

**UNIVERSITY OF CALIFORNIA, SAN FRANCISCO**
**CONSENT TO BE IN RESEARCH**

**Study Title:** Mapping the Intersection: Self-Identification and Genetic Ancestry

This is a research study, and you do not have to take part. You are being asked to take part in this study because you are registered with the National Marrow Donor Program (NMDP).

In this study, the researchers are doing a survey to learn more about the relationship between how people identify themselves with respect to race and ethnicity, and what their genetics tell us about their ancestry. The NMDP already has your genetic data for the genes involved in transplant, so there is no need to collect that information again. These data will be analyzed as part of this study.The National Human Genome Research Institute is paying for this research. About 50,000-100,000 people will participate in this study.

**What will happen if I take part in this study?**
If you agree to be in this study, you will complete a survey that begins on the next web page. The survey asks about your ancestry. It will take you about 5-10 minutes to complete the survey.

**Are there any risks to me or my privacy?**
Some of the survey questions may make you feel uncomfortable or raise unpleasant memories. You are free to skip any question.

We will do our best to protect the information we collect from you. Information that identifies you will be kept secure. The survey itself will not ask for details that directly identify you, such as your name or address. Please do not put this information on your survey. The completed surveys will be kept secure and separate from information that identifies you, and only a small number of researchers will have direct access to completed surveys. If this study is published or presented at scientific meetings, names and other information that might identify you will not be used.

**Are there benefits?**
There is no benefit to you. The survey results will be used for research.

**Can I say “No”?**
Yes, you do not have to complete the survey. If you choose not to be in this study you will remain registered as a potential bone marrow donor.

**Are there any payments or costs?**
You will not be paid for completing the survey. There are no costs to you.

**Who can answer my questions about the study?**
Answers to many questions may be found in the study frequently asked questions (FAQ): http://bethematch.org/HD/Ancestry-Study-FAQ/

You can talk with the study researcher about any questions not answered by the FAQ, concerns, or complaints you have about this study. Contact the study researcher Dr. Jill Hollenbach at 415-502-7289.

If you wish to ask questions about the study or your rights as a research participant to someone other than the researchers or if you wish to voice any problems or concerns you may have about the study, please call the Office of the Committee on Human Research at 415-476-1814.

**CONSENT**
PARTICIPATION IN RESEARCH IS VOLUNTARY.

You can print a copy of this consent form to keep for your records.

**If you wish to be in this study, please click “Continue.”**

- ☐ CONTINUE
- ☐ NO- I do not wish to participate in this study

Thank you for agreeing to participate in this important research study!

While taking the survey, *please do not use the back and forward buttons on your browser.* Once you have finished answering each question, click the button with the two arrows shown in the lower right hand corner of your screen to move on to the next question. You will not be able to return to previous questions during the survey, so please read each question carefully before you respond.

We appreciate your time and attention.

### Race and Ethnicity

*What is your race?*

Mark one or more boxes to show the racial or ethnic group(s) you use to describe yourself.

- ☐ American Indian or Alaska Native
- ☐ Asian
- ☐ Black or African American
- ☐ Hispanic or Latino
- ☐ Native Hawaiian or other Pacific Islander
- ☐ White
- ☐ Other (please specify)

*How do other people in this country typically classify you?*

Mark <u>one</u> selection to show the racial or ethnic group most Americans would use to describe you.

- » American Indian or Alaska Native
- » Asian
- » Black or African American
- » Hispanic or Latino
- » Native Hawaiian or other Pacific Islander
- » White
- » Other (please specify)

### Ancestry

*Were you born in the United States?*

- Yes
- No

*Where were your parents and grandparents born?*

Please include a response for each parent and grandparent.

| | Born in the U.S. | Born outside the U.S. | Don't know |
|---|---|---|---|
| Father | | | |
| Mother | | | |
| Paternal grandfather (your father's father) | | | |
| Paternal grandmother (your father's mother) | | | |
| Maternal grandfather (your mother's father) | | | |
| Maternal grandmother (your mother's mother) | | | |

*From what countries or parts of the world did your ancestors come?*

Select as many categories from the list below as needed to fully describe the origins of your family. If all of your grandparents were born in the United States, answer based on where your ancestors came from before they arrived in North America.

- [ ] Western Europe (England, Ireland, France, Germany, The Netherlands, etc.)
- [ ] Southern Europe (Italy, Spain, Turkey, etc.)
- [ ] Eastern Europe (Czech Republic, Poland, Russia, etc.)
- [ ] Scandinavia (Denmark, Norway, Sweden, etc.)
- [ ] East Asia (China, Japan, Korea, etc.)
- [ ] South Asia (India, Pakistan, Sri Lanka, etc.)
- [ ] Southeast Asia (Indonesia, Philippines, Vietnam, etc.)
- [ ] Pacific Islands (Hawaii, Guam, Samoa, etc.)
- [ ] Caribbean (Cuba, Puerto Rico, Trinidad and Tobago, etc.)
- [ ] Central or South America (Mexico, Nicaragua, Peru, etc.)
- [ ] Middle East (Iran, Lebanon, Saudi Arabia, etc.)
- [ ] Northern Africa (Egypt, Libya, Morocco, etc.)
- [ ] Sub-Saharan Africa (Kenya, Nigeria, Zimbabwe, etc.)
- [ ] American Indian (Navajo, Mayan, Tlingit, etc.)
- [ ] African American
- [ ] I do not know some, or all, of my family origins

Suppose you could describe a person using 100 points to represent all of their ancestries. For example, if you thought someone was mostly African but also part Latino, you might allocate 90 points for Sub-Saharan African and 10 points for Central or South American.

*How would you describe your ancestry using this 100-point system?*

Please indicate the number of points from 1-100 for each of the family origins shown below to represent its relative contributions to your ancestry.

| | |
|---|---|
| Western Europe (England, Ireland, France, Germany, The Netherlands, etc.) | 0 |
| Southern Europe (Italy, Spain, Turkey, etc.) | 0 |
| Eastern Europe (Czech Republic, Poland, Russia, etc.) | 0 |
| Scandinavia (Denmark, Norway, Sweden, etc.) | 0 |
| East Asia (China, Japan, Korea, etc.) | 0 |
| South Asia (India, Pakistan, Sri Lanka, etc.) | 0 |
| Southeast Asia (Indonesia, Philippines, Vietnam, etc.) | 0 |
| Pacific Islands (Hawaii, Guam, Samoa, etc.) | 0 |
| Caribbean (Cuba, Puerto Rico, Trinidad and Tobago, etc.) | 0 |
| Central or South America (Mexico, Nicaragua, Peru, etc.) | 0 |
| Middle East (Iran, Lebanon, Saudi Arabia, etc.) | 0 |
| Northern Africa (Egypt, Libya, Morocco, etc.) | 0 |
| Sub-Saharan Africa (Kenya, Nigeria, Zimbabwe, etc.) | 0 |
| American Indian (Navajo, Mayan, Tlingit, etc.) | 0 |
| African American | 0 |
| Unknown | 0 |
| **Total** | 0 |

Many Americans can trace their ancestry to several different parts of the world. To better understand your family history, we would like to ask about the ancestry of a few specific biological relatives.

*Do you know the origin(s) or ancestry of one or more of your grandparents?*

Please select all that apply from the list below.

- ☐ Yes, my paternal grandfather (biological father's biological father)
- ☐ Yes, my paternal grandmother (biological father's biological mother)
- ☐ Yes, my maternal grandfather (biological mother's biological father)

- Yes, my maternal grandmother (biological mother's biological mother)
- No, I do not know the ancestry of any of my biological grandparents

*What ancestry or origin(s) best describe your paternal grandfather?*

- Western Europe
  (England, Ireland, France, Germany, The Netherlands, etc.)
- Southern Europe
  (Italy, Spain, Turkey, etc.)
- Eastern Europe
  (Czech Republic, Poland, Russia, etc.)
- Scandinavia
  (Denmark, Norway, Sweden, etc.)
- East Asia
  (China, Japan, Korea, etc.)
- South Asia
  (India, Pakistan, Sri Lanka, etc.)
- Southeast Asia
  (Indonesia, Philippines, Vietnam, etc.)
- Pacific Islands
  (Hawaii, Guam, Samoa, etc.)
- Caribbean
  (Cuba, Puerto Rico, Trinidad and Tobago, etc.)
- Central or South America
  (Mexico, Nicaragua, Peru, etc.)
- Middle East
  (Iran, Lebanon, Saudi Arabia, etc.)
- Northern Africa
  (Egypt, Libya, Morocco, etc.)
- Sub-Saharan Africa
  (Kenya, Nigeria, Zimbabwe, etc.)
- American Indian
  (Navajo, Mayan, Tlingit, etc.)
- African American
- I do not know some of my paternal grandfather's origins

*What ancestry or origin(s) best describe your paternal grandmother?*

- Western Europe
  (England, Ireland, France, Germany, The Netherlands, etc.)

- Southern Europe
  (Italy, Spain, Turkey, etc.)
- Eastern Europe
  (Czech Republic, Poland, Russia, etc.)
- Scandinavia
  (Denmark, Norway, Sweden, etc.)
- East Asia
  (China, Japan, Korea, etc.)
- South Asia
  (India, Pakistan, Sri Lanka, etc.)
- Southeast Asia
  (Indonesia, Philippines, Vietnam, etc.)
- Pacific Islands
  (Hawaii, Guam, Samoa, etc.)
- Caribbean
  (Cuba, Puerto Rico, Trinidad and Tobago, etc.)
- Central or South America
  (Mexico, Nicaragua, Peru, etc.)
- Middle East
  (Iran, Lebanon, Saudi Arabia, etc.)
- Northern Africa
  (Egypt, Libya, Morocco, etc.)
- Sub-Saharan Africa
  (Kenya, Nigeria, Zimbabwe, etc.)
- American Indian
  (Navajo, Mayan, Tlingit, etc.)
- African American
- I do not know some of my paternal grandmother's origins

*What ancestry or origin(s) best describe your maternal grandfather?*

- Western Europe
  (England, Ireland, France, Germany, The Netherlands, etc.)
- Southern Europe
  (Italy, Spain, Turkey, etc.)
- Eastern Europe
  (Czech Republic, Poland, Russia, etc.)
- Scandinavia
  (Denmark, Norway, Sweden, etc.)
- East Asia
  (China, Japan, Korea, etc.)
- South Asia

(India, Pakistan, Sri Lanka, etc.)

- Southeast Asia
(Indonesia, Philippines, Vietnam, etc.)
- Pacific Islands
(Hawaii, Guam, Samoa, etc.)
- Caribbean
(Cuba, Puerto Rico, Trinidad and Tobago, etc.)
- Central or South America
(Mexico, Nicaragua, Peru, etc.)
- Middle East
(Iran, Lebanon, Saudi Arabia, etc.)
- Northern Africa
(Egypt, Libya, Morocco, etc.)
- Sub-Saharan Africa
(Kenya, Nigeria, Zimbabwe, etc.)
- American Indian
(Navajo, Mayan, Tlingit, etc.)
- African American
- I do not know some of my maternal grandfather's origins

*What ancestry or origin(s) best describe your maternal grandmother?*

- Western Europe
(England, Ireland, France, Germany, The Netherlands, etc.)
- Southern Europe
(Italy, Spain, Turkey, etc.)
- Eastern Europe
(Czech Republic, Poland, Russia, etc.)
- Scandinavia
(Denmark, Norway, Sweden, etc.)
- East Asia
(China, Japan, Korea, etc.)
- South Asia
(India, Pakistan, Sri Lanka, etc.)
- Southeast Asia
(Indonesia, Philippines, Vietnam, etc.)
- Pacific Islands
(Hawaii, Guam, Samoa, etc.)
- Caribbean
(Cuba, Puerto Rico, Trinidad and Tobago, etc.)
- Central or South America
(Mexico, Nicaragua, Peru, etc.)

- Middle East
  (Iran, Lebanon, Saudi Arabia, etc.)
- Northern Africa
  (Egypt, Libya, Morocco, etc.)
- Sub-Saharan Africa
  (Kenya, Nigeria, Zimbabwe, etc.)
- American Indian
  (Navajo, Mayan, Tlingit, etc.)
- African American
- I do not know some of my maternal grandmother's origins

*Do you know the origin(s) or ancestry of either of your biological parents?*

Please select all that apply from the list below.

- Yes, my biological mother
- Yes, my biological father
- I do not know the ancestry of either of my biological parents

*What ancestry or origin(s) best describe your biological mother?*

- Western Europe
  (England, Ireland, France, Germany, The Netherlands, etc.)
- Southern Europe
  (Italy, Spain, Turkey, etc.)
- Eastern Europe
  (Czech Republic, Poland, Russia, etc.)
- Scandinavia
  (Denmark, Norway, Sweden, etc.)
- East Asia
  (China, Japan, Korea, etc.)
- South Asia
  (India, Pakistan, Sri Lanka, etc.)
- Southeast Asia
  (Indonesia, Philippines, Vietnam, etc.)
- Pacific Islands
  (Hawaii, Guam, Samoa, etc.)
- Caribbean
  (Cuba, Puerto Rico, Trinidad and Tobago, etc.)
- Central or South America

(Mexico, Nicaragua, Peru, etc.)

- Middle East
  (Iran, Lebanon, Saudi Arabia, etc.)
- Northern Africa
  (Egypt, Libya, Morocco, etc.)
- Sub-Saharan Africa
  (Kenya, Nigeria, Zimbabwe, etc.)
- American Indian
  (Navajo, Mayan, Tlingit, etc.)
- African American
- I do not know some of my biological mother's origins

*What ancestry or origin(s) best describe your biological father?*

- Western Europe
  (England, Ireland, France, Germany, The Netherlands, etc.)
- Southern Europe
  (Italy, Spain, Turkey, etc.)
- Eastern Europe
  (Czech Republic, Poland, Russia, etc.)
- Scandinavia
  (Denmark, Norway, Sweden, etc.)
- East Asia
  (China, Japan, Korea, etc.)
- South Asia
  (India, Pakistan, Sri Lanka, etc.)
- Southeast Asia
  (Indonesia, Philippines, Vietnam, etc.)
- Pacific Islands
  (Hawaii, Guam, Samoa, etc.)
- Caribbean
  (Cuba, Puerto Rico, Trinidad and Tobago, etc.)
- Central or South America
  (Mexico, Nicaragua, Peru, etc.)
- Middle East
  (Iran, Lebanon, Saudi Arabia, etc.)
- Northern Africa
  (Egypt, Libya, Morocco, etc.)
- Sub-Saharan Africa
  (Kenya, Nigeria, Zimbabwe, etc.)
- American Indian
  (Navajo, Mayan, Tlingit, etc.)

- African American
- I do not know some of my biological father's origins

### Knowledge check

*How much would you say you know about your family history on your biological mother's side?*

- Nothing at all
- A little
- A lot

*How much would you say you know about your family history on your biological father's side?*

- » Nothing at all
- » A little
- » A lot

*Have you ever done any of the following to seek out information about your ancestry or family history?*

Please select all that apply from the list below.

- Asked family members questions about family history
- Gone through family documents to find information
- Used a genealogy website, such as ancestry.com, familysearch.org
- Sent away for birth, death or marriage certificates or other official documents
- Gone to a library or archive to look for family records
- Taken a genetic ancestry test
- Other (please specify)
- None of these

### Confidence family history

Some families do not talk about or are unaware of all their ancestral origins. On a scale from 1 (extremely unlikely) to 5 (extremely likely), *how likely do you think it is that you have any of the following ancestries:*

| | Extremely unlikely | Very unlikely | Neither likely nor unlikely | Very likely | Extremely likely |
|---|---|---|---|---|---|
| African | ◯ | ◯ | ◯ | ◯ | ◯ |
| American Indian | ◯ | ◯ | ◯ | ◯ | ◯ |
| East Asian | ◯ | ◯ | ◯ | ◯ | ◯ |
| South Asian | ◯ | ◯ | ◯ | ◯ | ◯ |
| Jewish | ◯ | ◯ | ◯ | ◯ | ◯ |
| Middle Eastern | ◯ | ◯ | ◯ | ◯ | ◯ |
| Scandinavian | ◯ | ◯ | ◯ | ◯ | ◯ |
| Southern European | ◯ | ◯ | ◯ | ◯ | ◯ |
| Eastern European | ◯ | ◯ | ◯ | ◯ | ◯ |
| Western European | ◯ | ◯ | ◯ | ◯ | ◯ |

### Ancestry testing

*How familiar are you with genetic ancestry testing? Would you say you are...*

- ◯ Very familiar
- ◯ Somewhat familiar
- ◯ Somewhat unfamiliar
- ◯ Very unfamiliar

*If you were offered a free genetic ancestry test, would you be interested in taking it?*

- ◯ Yes
- ◯ No

Please share with us the reason(s) why you would not be interested in taking a genetic ancestry test.

*If you took a genetic ancestry test and were told your ancestors were from one or more of these origins, how happy would you be?*

For each ancestry, please indicate your level of happiness, from very unhappy to very happy.

| | Very unhappy | Unhappy | Neither happy nor unhappy | Happy | Very happy |
|---|---|---|---|---|---|
| » African | ○ | ○ | ○ | ○ | ○ |
| » American Indian | ○ | ○ | ○ | ○ | ○ |
| » East Asian | ○ | ○ | ○ | ○ | ○ |
| » South Asian | ○ | ○ | ○ | ○ | ○ |
| » Jewish | ○ | ○ | ○ | ○ | ○ |
| » Middle Eastern | ○ | ○ | ○ | ○ | ○ |
| » Scandinavian | ○ | ○ | ○ | ○ | ○ |
| » Southern European | ○ | ○ | ○ | ○ | ○ |
| » Eastern European | ○ | ○ | ○ | ○ | ○ |
| » Western European | ○ | ○ | ○ | ○ | ○ |

### Basic Demographics

*Around the time you were 16, were you living with both your biological mother and father?*

- ○ Yes, both
- ○ No, biological mother only
- ○ No, biological father only
- ○ No, neither

*What was the highest educational degree your mother received?*

- ○ Did not finish high school

- High school degree or equivalent
- Associate's degree
- Bachelor's degree
- Graduate or professional degree
- Don't know

*What was the highest educational degree your father received?*

- » Did not finish high school
- » High school degree or equivalent
- » Associate's degree
- » Bachelor's degree
- » Graduate or professional degree
- » Don't know

*What is the highest educational degree you have received?*

- Did not finish high school
- High school degree or equivalent
- Associate's degree
- Bachelor's degree
- Graduate or professional degree

*Please select the state where you lived around the time you were 16.*

*Please select the state where you currently live.*

*What is your age?*

- 18 to 24 years
- 25 to 34 years
- 35 to 44 years
- 45 to 54 years
- 55 to 64 years
- Age 65 or older

### Open-ended

*Is there any information about your race or ancestry that you would like to share with us, that was not captured by the previous questions or answer options?*

Please provide any additional details that you think would be important to fully understand your family origins and ancestry.
